# Automated non-invasive laser speckle imaging of the chick heart rate and extraembryonic blood vessels and their response to nifedipine and amlodipine drugs

**DOI:** 10.1101/2024.08.26.609812

**Authors:** Carol Readhead, Simon Mahler, Zhenyu Dong, Yuki Sato, Changhuei Yang, Marianne E. Bronner

**Author notes:** These authors contributed equally (Carol Readhead, Simon Mahler).

## Abstract

Using our recently developed laser speckle contrast imaging (LSCI) to visualize blood vessels and monitor blood flow, here we test the utility of the chick embryo for drug screening. To this end, we examined the effects of antihypertensive agents Nifedipine and Amlodipine, belonging to the L-type calcium channel antagonist family, on blood flow visualized noninvasively through the intact shell. Guided by the live view mode, the drugs were injected through the shell and ventral to HH16-19 chick embryos. Our results show a significant reduction in the chick heart rate, blood flow, and vascular size within 5-20 minutes after Nifedipine or Amlodipine injection. For moderate Nifedipine concentrations, these parameters returned to initial values within 2-3 hours. In contrast, Amlodipine showed a rapid reduction in heart rate and blood flow dynamics at a more than ten times higher concentration than Nifedipine. These findings show that our LSCI system can monitor and distinguish the chick heart’s response to injected drugs from the same family. This serves as proof-of-concept, paving the way for a rapid, cost effective, and quantitative test system for screening drugs that affect the cardiovascular system of live chick embryos. Live noninvasive imaging may also provide insights into the development and functioning of the vertebrate heart.

**Graphical Abstract:** 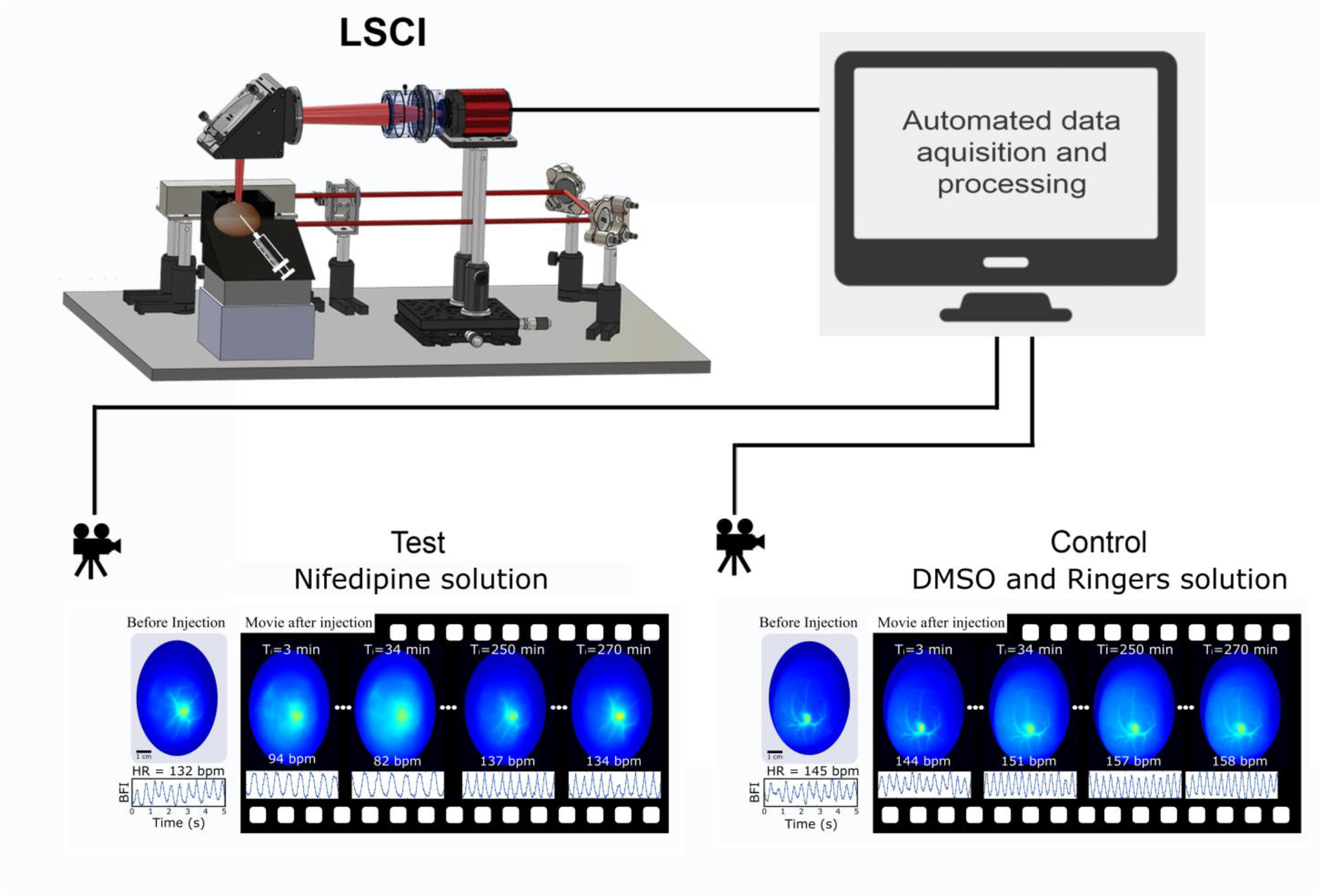

**Highlights:**

- Non-invasive Laser Speckle Contrast Imaging (LSCI) of the chick chorioallantoic membrane (CAM) in whole incubated eggs
- Simultaneous recording images of the CAM, dynamics of blood flow, and heart rate
- Live view mode to identify size, heart position, and location of the embryo in the egg
- Automated system for data acquisition and analysis
- Longitudinal quantification of the impact of a calcium channel antagonists, nifedipidine and amlodipine on the embryonic heart rate, CAM’s blood flow, size and number of vessels

## 1. Introduction

The domestic chicken, *Gallus gallus domesticus*, is valuable not only for the study of embryonic development but also as a model system for biomedical research (Ribatti et al., 2021; Ribatti and Annese, 2023). During a chick embryo’s development, a rich nextwork of blood vessels grow in the extraembryonic membranes that provide a means for gas and waste exchange. The chick embryo follows a specific developmental process categorized into HH-stages by Hamburger and Hamilton, with each stage corresponding to a distinct developmental feature of the embryo (Hamburger and Hamilton, 1951). At stage HH9, the highly vascularized chorionic and allantoic membrances fuse to form the chorioallantoic membrane (CAM) (DeFouw et al., 1989). At stage HH10 (approximately 33 hours), the heart starts to beat (Kamino, 1989) starting the blood circulation (Al Naieb et al., 2013; Martinsen, 2005). This accessible embryonic cardiovascular system has been used as a model to study: heart development, angiogenesis, anti-hypertensive drugs, metastastatic tumors and organoids (Burggren and Rojas Antich, 2020; Gandhi et al., 2020; Goenezen et al., 2016; Nowak-Sliwinska et al., 2014; Ribatti, 2016). Moreover, the highly vascularized chick extraembryonic blood vessels, including those of the CAM have been used to explore the effects of heart inhibitors (As et al., 2018; Mei et al., 2001).

Developmental and biomedical studies are usually done *in ovo* or *ex ovo*. For in ovo studies a window is cut in the shell to expose the embryo for mechanical or molecular manipulation. The window is then covered with transparent tape and the egg is incubated (Bronner-Fraser, 1996; Sukumaran et al., 2023). Alternatively, the embryo and extra blood vessels can be isolated and grown *in vitro* (New, 1955). When direct access to the embryo is not required, non-invasive imaging of the chick embryonic cardiovascularsystem is beneficial as opening the egg disrupts the natural development of the embryo and diminishes the overall survival rate of the embryo (Jia et al., 2023; Khaliduzzaman, 2022).

Hypertension is a leading risk factor for cardiovascular disease and premature death. With more than 30% of adults worldwide affected by hypertension (Mills et al., 2020; Stewart, 2023), extensive studies have been done on comparing different classes of treatments (Reinhart et al., 2023). An important class of drugs are the antihypertensive drugs belonging to the L-type calcium channel antagonist family, which include Nifedipine and Amlodipine (Fleckenstein, 1983). Nifedipine (Mills et al., 2020) is sometimes used in conjunction with herbal remedies (Wang et al., 2022), while Amlodipine is commonly used for treatment due to it long half-life (Mostafa et al., 2022).

Recently, we have developed a non-invasive imaging system that overcomes the opacity of the shell and visualizes the beating heart of the chick embryo and the flow of blood in the the extraembryonic vessels of the CAM (Dong et al., 2024). The system is based on laser speckle contrast imaging (LSCI) that can visualizes blood vessels and monitor blood flow in a variety of tissues(Draijer et al., 2009; Dunn et al., 2001; Mahler et al., 2023; Senarathna et al., 2013). In our LSCI system, the beating heart and blood flow index can be viewed in realtime (Dong et al., 2024). For longitudinal studies of the cardiovascular system, we refined the LSCI setup to include an incubator with automated data acquisition and analysis at chosen time points.

In this paper, we show that our LSCI system with an integrated incubator, developed for live imaging of the chick heart and extraembryonic blood flow, can be used in longitudinal studies of the embryonic heart and the drugs and factors that can affect its function. Such LSCI system is non-invasive, and does not alter the natural development of the chick embryo. As proof-of-concept, we tested two drugs that are commonly used to treat hypertension in humans: Nifedipine and Amlodipine. These were injected through the shell ventral to the embryo at HH16-19. The needle was guided by the LSCI live view mode. Our results show a significant reduction in the chick heart rate, blood flow, and vascular size within 5 to 15 minutes after Nifedipine or Amlodipine injection. For moderate Nifedipine concentrations, these parameters returned to initial values within 2-3 hours. In contrast, Amlodipine showed a rapid reduction in heart rate and blood flow dynamics at more than ten times higher concentration than Nifedipine. Thus, the LSCI system can monitor and distinguish the chick heart’s response to injected drugs from the same family. This serves as proof-of-concept, paving the way for a rapid, cost effective, and quantitative test system for screening drugs that affect the cardiovascular functioning of live chick embryos.

## 2. Material and Methods

### 2.1 Laser Speckle Contrast Imaging (LSCI)

The blood vessel network of the CAM was imaged using laser speckle contrast imaging (LSCI). This technique relies on laser speckles generated by the scattering of coherent laser light in the sample (Boas and Dunn, 2010; Chriki et al., 2018; Draijer et al., 2009; Dunn et al., 2001; Goodman, 2007; Mahler et al., 2020). There has been significant interest in laser speckles as a method for visualizing blood vessels and monitoring blood flow in many types of living tissues (Draijer et al., 2009; Dunn, 2012; Dunn et al., 2001; Mahler et al., 2023; Senarathna et al., 2013) including avian embryos (Chen et al., 2023; Du et al., 2020; Padmanaban et al., 2021; Yang et al., 2013).

Speckles arise from the interferences of the randomly scattered laser light by the sample and follow a specific temporal evolution. As components such as blood within the sample move, the speckle field experiences temporal changes. By imaging the speckle patterns with a camera and analyzing the fluctuations among the recorded camera images, it is possible to reconstruct an image of the blood flow distribution within the sample. This is achieved by calculating the speckle contrast from the recorded camera images, which is defined as the ratio between the standard deviation and the mean value of the pixel intensities. As the moving blood cells in the vessels exhibit greater movements, the speckle contrast is lower, and vice-versa. A more detailed description can be found in (Dong et al., 2024).

In our case, we used temporal LSCI (Li et al., 2006; Senarathna et al., 2013), where the camera recorded a sequence of N = 100 speckle frames images when the laser was on and another 100 noise frames when the laser was blocked with the same exposure time T to generate noise subtracted speckle frames 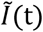 to reduce noise. The noise frames were averaged to a single frame and substracred from the speckle frames (Dong et al., 2024). The temporal speckle contrast *K*_*t*_ was calculated for each camera pixel 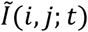 in the temporal domain over the N noise subtracted speckle frames as:

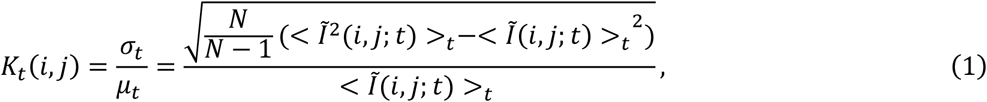

where 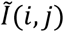 is the intensity at the pixel row *i* and column *j, t* is the time at which the speckle pattern was recorded by the camera, and <>_*t*_ indicates temporal averaging over time occurring at pixel *i, j*. The speckle contrast corresponds to the ratio between the standard deviation *σ*_*t*_ and the temporal mean *μ*_*t*_ of the N recorded speckle pattern images. The blood flow index (BFI) was calculated from the speckle contrast as (Huang et al., 2024b):

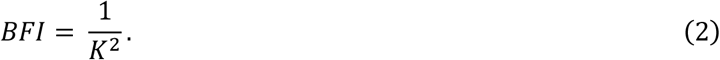

The blood flow index provide relative blood flow information, accounting for the total volume of blood moved in a given time period. Any alteration in the BFI means that there is a change in either the blood pressure or a change in the diameter of the blood vessel. Figure 1 shows a typical example of an egg’s blood vessel image and reconstructed blood flow dynamics using tLSCI. A clear visualization of the blood vessel network can be observed encompassing vessels of varying sizes, and the heart rate can be measure from the blood flow dynamics.

**Fig. 1.**
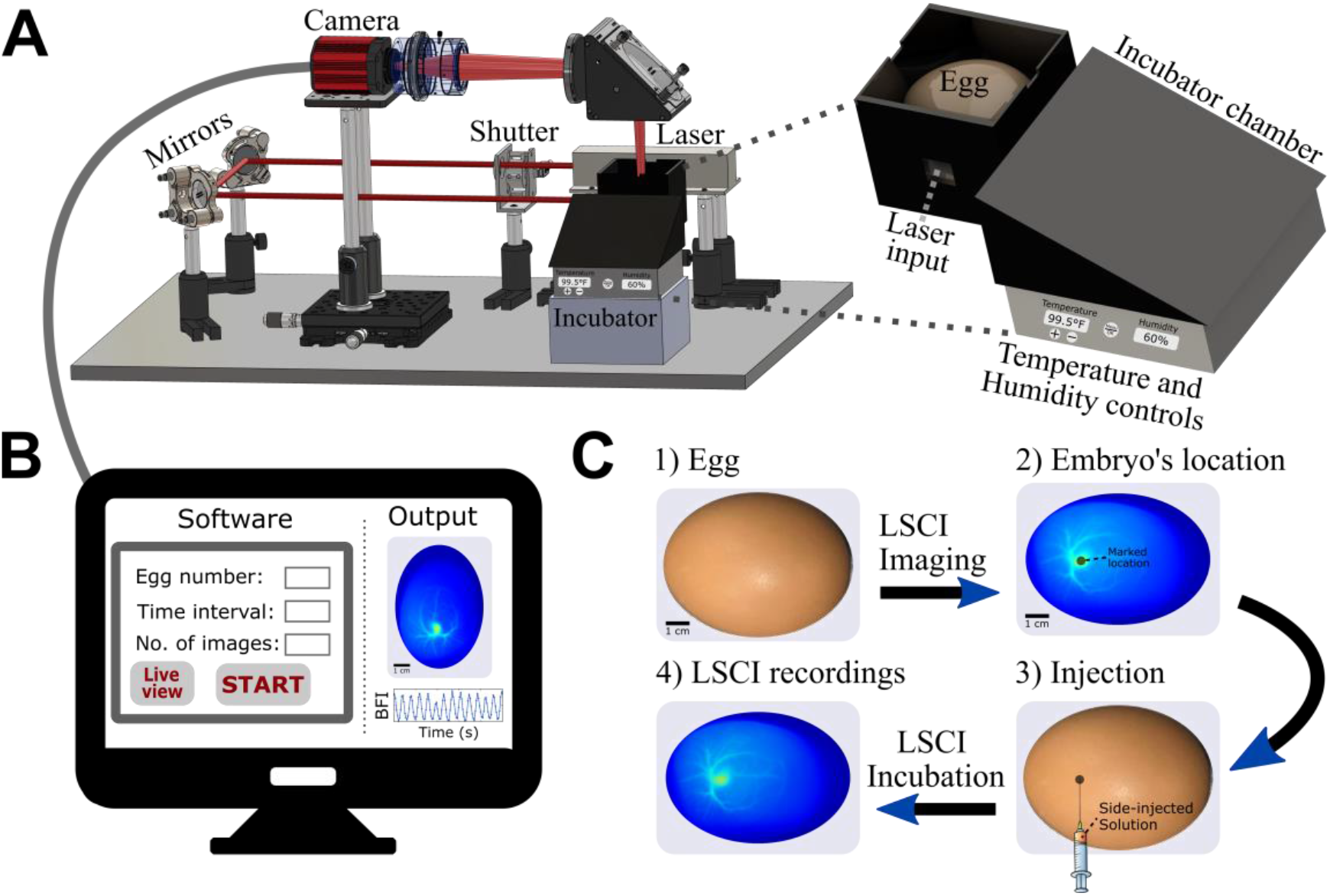
Integrated incubation in LSCI system for imaging chicken egg blood vessels post injection. (A) Schematic of the experimental arrangement of the laser speckle contrast imaging (LSCI) system incorporating an incubator. Right: detailed view of the LSCI incubator. (B) Software interface for automated monitoring of the chicken egg’s blood vessels at regular intervals. (C) Drug-injection protocol using LSCI imaging to determine the location of the embryo’s for accurate injection. The injection was done through the lateral side of the egg shell, so as to not damage the embryo.

### 2.2 Extracting the blood flow dynamics from LSCI recordings

We demonstrated previously that the blood flow dynamics in chicken eggs can be derived from LSCI recordings (Dong et al., 2024). Instead of using the N = 100 speckle frames, one can monitor blood flow dynamics by applying a sliding window of n = 3 adjacent speckle frames to calculate a time-varying temporal speckle contrast. This contrast can be converted into blood flow values over time. With our camera recording at 21 frames per second, blood flow can be recorded with a temporal resolution of 21 data points per second.

The blood flow extraction method was described in details in(Dong et al., 2024) and has been shown to provide results similar to those obtained with the speckle visibility spectroscopy (SVS) method (Huang et al., 2024b; Mahler et al., 2023), where both LSCI and SVS measured the same heart rate and blood flow dynamics of a chicken egg LSCI dataset. In SVS, blood flow is calculated from a single speckle frame using spatial speckle contrast, with each camera pixel acting as a sample point. For significant results with SVS, a camera frame rate of 40 fps or higher is required.

### 2.3 Experimental arrangement of LSCI integrated incubation

To study the effects of drug injection into the CAM of a chicken egg, we incorporated an incubator into our LSCI system, Fig. 1. The LSCI system is presented in Fig. 1A (left). The laser source used was a single frequency continuous wave 852 nm laser [Spectra-Physics DL852-300-SO] with a output power of 230 mW. Using a pair of reflective mirrors, the laser light side-illuminated the chicken egg (Dong et al., 2024). The scattered light exiting the egg was collected by an imaging system consisting of a mirror with an aperture to control the field-of-view and to filter out undesired stray light, and a camera with a 50 mm focal length lens [Edmund Optics #86-574]). The magnification of the imaging system was 0.2 and the numerical aperture was 0.03. The camera [Thorlabs CS126CU] was set with an exposure time of 10 ms and at a speed of 21 frames-per-second, had a resolution of 3000 × 4096 pixels, a pixel pitch size of 3.45 × 3.45 µm, and the speckle size was about two pixels per speckle. A digitally controlled optical shutter blocked the laser source when the LSCI system was not recording.

A detailed view of the incubator is shown in Fig. 1A (right). For our application, we replaced the top part of the incubator by a homemade 3D-printed chamber together with an egg holder designed to fit inside the LSCI system. The printing was done using Anycubic resin 3D printer. A 1.4 × 1.4 mm square aperture covered with transparent tape served as the laser input for side illuminating the egg. The surface above the egg was covered with a transparent plastic window. To verify that the homemade 3D-printed incubator chamber provided sufficient heat and humidity for natural egg development, we incubated five eggs from day 0 to day 5. All five eggs developed normally to the expected stage HH25 after 5 days of incubation.

The LSCI system was enclosed in a black posterboard box to prevent room and stray light noises. The LSCI recordings were conducted remotely using a Matlab graphical user interface (GUI), as shown in Fig. 1B. The GUI included a live view mode, offering the capability of viewing the chicken egg every second using three speckle frames to reconstruct the blood vessel image (Dong et al., 2024). Additionally, the GUI featured an automated LSCI recording function, allowing the user to set the recording interval (in minutes) and the number of images to capture. The GUI automated the optical shutter and camera during recording, requiring no further user intervention. The output from the GUI, displayed on the right side of Fig. 1B, consisted of an image of the egg’s blood vessels and the blood flow dynamics over a period of 5 seconds.

### 2.4 Injecting the egg

Figure 1C shows the protocol for injecting a solution into an egg. First, an incubated egg was selected for injection. Second, using the LSCI live view mode, the position of the embryo’s heart was ascertained and marked on the eggshell. Third, at the marked location, a 22-gauge curved teasing needle was used to pierce a hole into the lateral side of the eggshell. A syringe needle of 23-gauge was brought to the embryo marker location using the LSCI live view mode. This resulted in a hole into the eggshell with diameter between 0.6 to 0.8 mm. We believe that by using a higher gauge needles would result in a smaller hole, less than 0.5 mm diameter. The hole was covered by transparent tape, ensuring a continued development of the embryo. The marked location was sanitized before, and after the injection with alcohol wipes. After injection, the egg was placed in the LSCI incubator and imaged at regular interval using the GUI of Fig. 1B.

### 2.5 Preparation of the Nifedipine and Amlodipine solutions

The impact of Nifedipine and Amlodipine (Sigma-Aldrich), two L-Type Ca^2a^ channel antagonists were tested on the chick embryonic heart. The drugs were injected into the egg at the ventral position of the embryo. Nifedipine, being less soluble was first dissolved in DMSO at a concentration of 50mg/ml according to the manufactures instructions, and was kept shielded from UV light (Ali, 1990; Bottorff et al., 1984; Martindale and Reynolds, 1993). Each day of the experiment a fresh stock solution was made and then diluted in sterile Howard Ringers (Ringers). About 1.5 nmol, 3 nmol or 5 nmol of Nifedipine (molecular weight of 346 g/mol) were injected in a volume of 25µL, 50µL or 100µL respectively. The same dilution of DMSO + Ringers was used as a control solution (1 µL of DMSO in 2,500 µL Ringers). Studies in an electrolyte solution indicated that the Nifedipine’s stability declined to about 90% of its orginal value within six hours after preparation (Bottorff et al., 1984) which was also true for these experiments. When dissolved, Nifedipine is also unstable over time and extremely photosensitive, especially to UV light (Ali, 1990; Bottorff et al., 1984; Martindale and Reynolds, 1993).

A quantity of 5 mg of Amlodipine (molecular weight of 567 g/mol) was diluted in 10 mL of lukewarm sterile Ringers. It was effective at much higher doses than Nifedipine and was injected at about 45 nmol or 65 nmol in a volume of 50µL or 75µL respectively. Diluted Amlodipine was stable for several days at room temperature and was not affected by UV light exposition.

### 2.6 Eggs supply and use

In all our experiments, we used brown fertile eggs obtained from Sun Valley farms (CA). They were incubated in Hethya, HHD, and Kebonnixs commercial incubators. These incubators automatically regulated the temperature within 99-101°F range and gently turned the eggs every 60 or 90 min. The humidity in the incubators was maintained within the 50-70% range. After 3-4 days of incubation the embryos that had developed in a stage range between HH16-19 were used for imaging. The animal research for this study received confirmation from the Caltech Office of Laboratory Animal Resources. In all our studies, chicken eggs were incubated no later than day 12. The disposal of the chicken egg embryo was performed in compliance with the Caltech Institutional Animal Care and Use Committee policy (IACUC).

## 3. Results

### 3.1 Testing and validating the LSCI system with Nifedipine

To validate this system for longitundinal drug studies, as a-proof-of-concept, we first tested Nifedipine. For that, we injected and recorded two different eggs using our LSCI incubation system. The results are presented in Fig. 2. One egg corresponding to the control group was injected by 50 µL of a solution consisting of 1 µL DMSO and 2.5 mL Ringer’s. The second egg corresponding to the testing group was injected by 50 µL of Nifedipine + DMSO solution, prepared by dissolving 50 mg of Nifedipine in 1 mL of DMSO and diluted in Ringers. As shown in Fig. 2A, there was little change in the heart rate or the pattern of the CAM blood vessels for the control egg over the duration of the recording (270 min). The embryo exhibited almost no change in the heart rate or the pattern of blood vessels over the time of the experiment (250 minutes)(Fig. 2B).

**Fig. 2.**
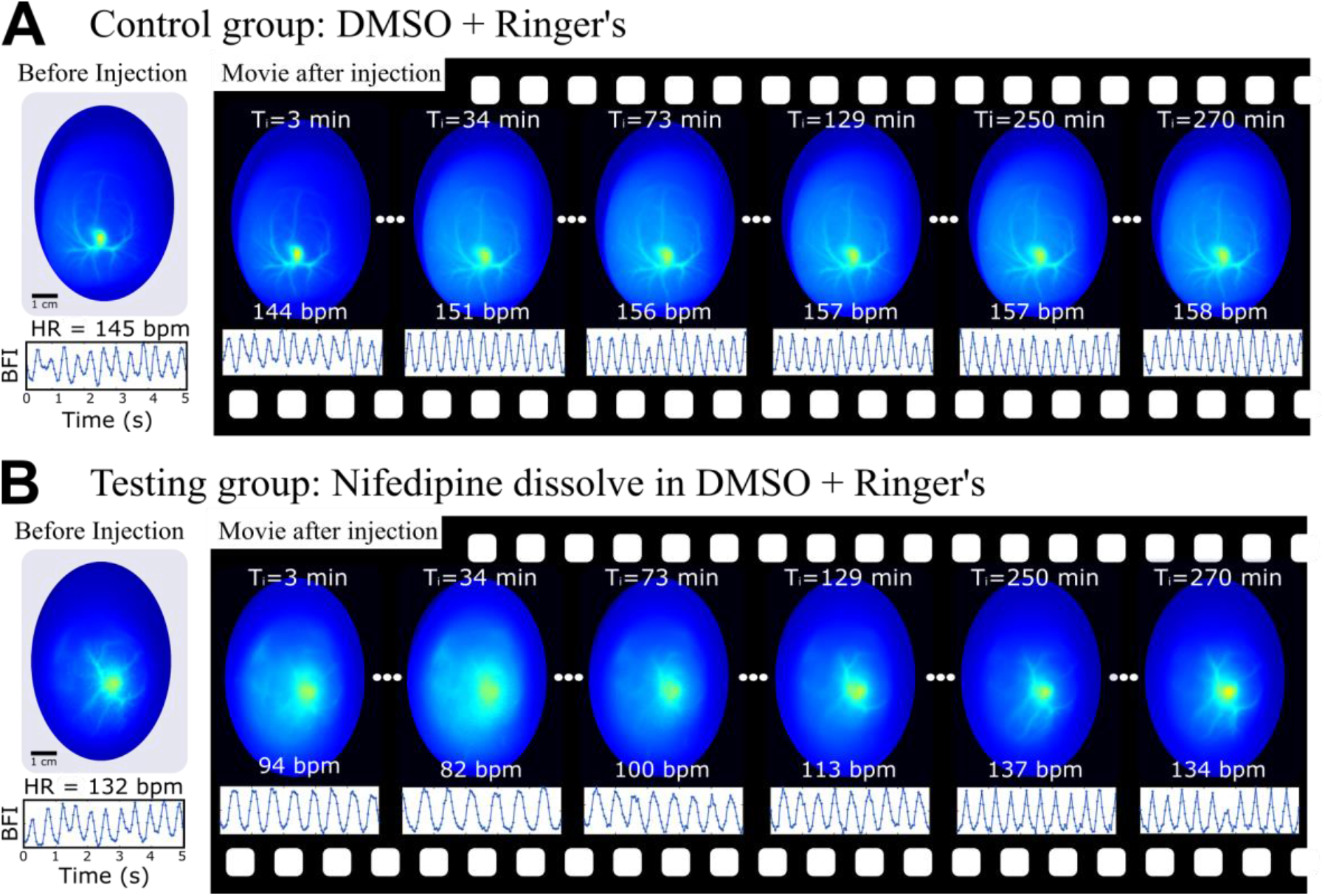
Monitoring the blood vessel network and blood flow dynamics of chicken eggs after injecting a solution. (A) Control group: egg injected with 50 µL of a solution consisting of 1 µL DMSO and 2.5 mL Ringer’s. (B) Testing group: egg injected with 50 µL of a solution consisting of 3 nmole of Nifedipine. This was prepared by first dissolving 50 mg/mL of Nifedipine in DMSO and diluting 1 µL of this solution in 2.5 mL of Ringers. As shown, the heart rate and vascular structure of the control egg remained consistent across all recordings, while the Nifedipine-injected egg exhibited an initial reduction of blood flow in the CAM which resulted in loss of the vascular structure in the image. This was followed by revival of the heart rate, blood flow, and image of the vascular structure.

For the test egg, Nifedipine was injected. The heart rate dropped from 132 beat-per-minute (bpm) to 94 bpm and the CAM’s blood vessels were no longer clearly visible due to the decreased blood flow (Fig. 2B). An hour after injection, the heart rate increased to 100 bpm and by 250 minutes it had returned to the original pre-injection levels. The blood vessels were again clearly visible. These results are in agreement with previous observations on the action of Nifedipine on chick embryonic hearts *in vitro* (Mei et al., 2001). For each group, we repeated the tests on five different eggs, and observing similar behavior than in Fig. 2 for each group. We then re-incubated the eggs until day-12, when they were opened and dissected to assess their developmental stage and health. All five eggs in the control group developed naturally, showing no signs of malformation or illness during dissection. In contrast, all five eggs in the test group were found to be dead at the time of opening, with some having died at a later stage than others.

### 3.2 Extraction of features from blood flow images

To quantify and characterize the effects of injecting a drug solution onto the CAM, we extracted four features from the blood vessels images and blood flow graphs. These features are shown in Fig. 3. Figure 3A shows a typical blood vessel image obtained from the LSCI. Figure 3B presents the corresponding LSCI+ image, which segments out large blood vessels from LSCI images and identifies the blood vessels skeletons (black lines), the branch points (green dots), and CAM area (beige shaded area). The LSCI+ method consist of 12 steps, see (Dong et al., 2024) for details. Figure 3C (left) shows a typical blood flow dynamics of a chicken egg, obtained from LSCI (see Materials and Methods for more details). Figure 3C (right) shows the Fourier transform of the blood flow signal, where the heart rate can be measured (Huang et al., 2024b, 2024a; Mahler et al., 2023). Our current LSCI+ method is unable to segment smaller blood vessels or those with signals near background noise levels. However, we believe these issues can be addressed by developing an enhanced LSCI++ segmentation algorithm that incorporates image filters and multiple segmentation layers—one for large vessels and another for smaller vessels.

**Fig. 3.**
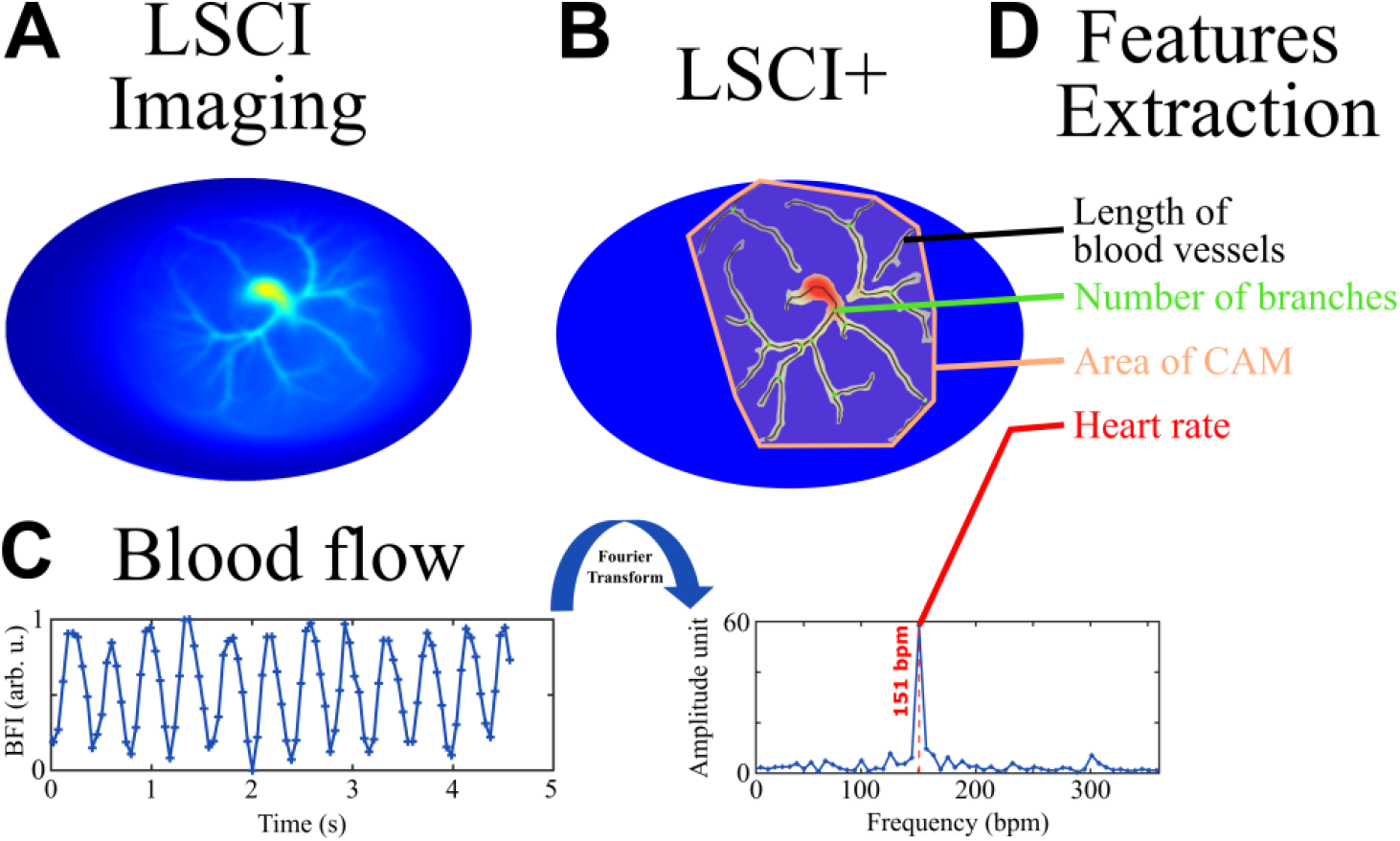
Features extraction from a LSCI recording of the blood vessels in a chicken egg. (A) Typical LSCI image of the blood vessels in a chicken egg. (B) An enhanced version of LSCI, called LSCI+, depicting blood vessel skeletons, branches, and the size of the CAM. (C) (left) Blood flow of the imaged chicken egg with. (right) Fourier frequencies obtained by Fourier Transforming the blood flow graph. The red dashed line shows the measured heart rate. (D) Extracted features from the LSCI recording.

Figure 3D presents the extracted features from LSCI+ and heart rate measurements. The length of the blood vessels was calculated as the total length of the vessel skeleton in the LSCI+ image. The number of branches was determined by counting the branch points connecting vessels and isolated vessels regions in the LSCI+ image. The vessel area was obtained by calculating the area of the CAM in the LSCI+ image. Finally, the heart rate was measured from the Fourier transform of the blood flow signal.

### 3.3 Quantification of the heart rate (bpm), length of blood vessels, number of branches, and area of CAM after injection of Nifedipine or Amlodipine

Using the LSCI incubation system and feature extraction method, we quantified the heart rate, length of blood vessels, number of branches, and area of the CAM after injecting Nifedipine (Fig.4A-C) or Amlodipine (Fig.5 A-C). For Nifedipine, each egg was injected with a different amount of Nifedipine. The first datapoint with negative time in Figs. 4A-C, corresponds to the features measured before injection. The visible length of the blood vessels (4B), the number of branches (4C) and the area of the CAM (4D) due to the the decrease in the blood flow index.

**Fig. 4.**
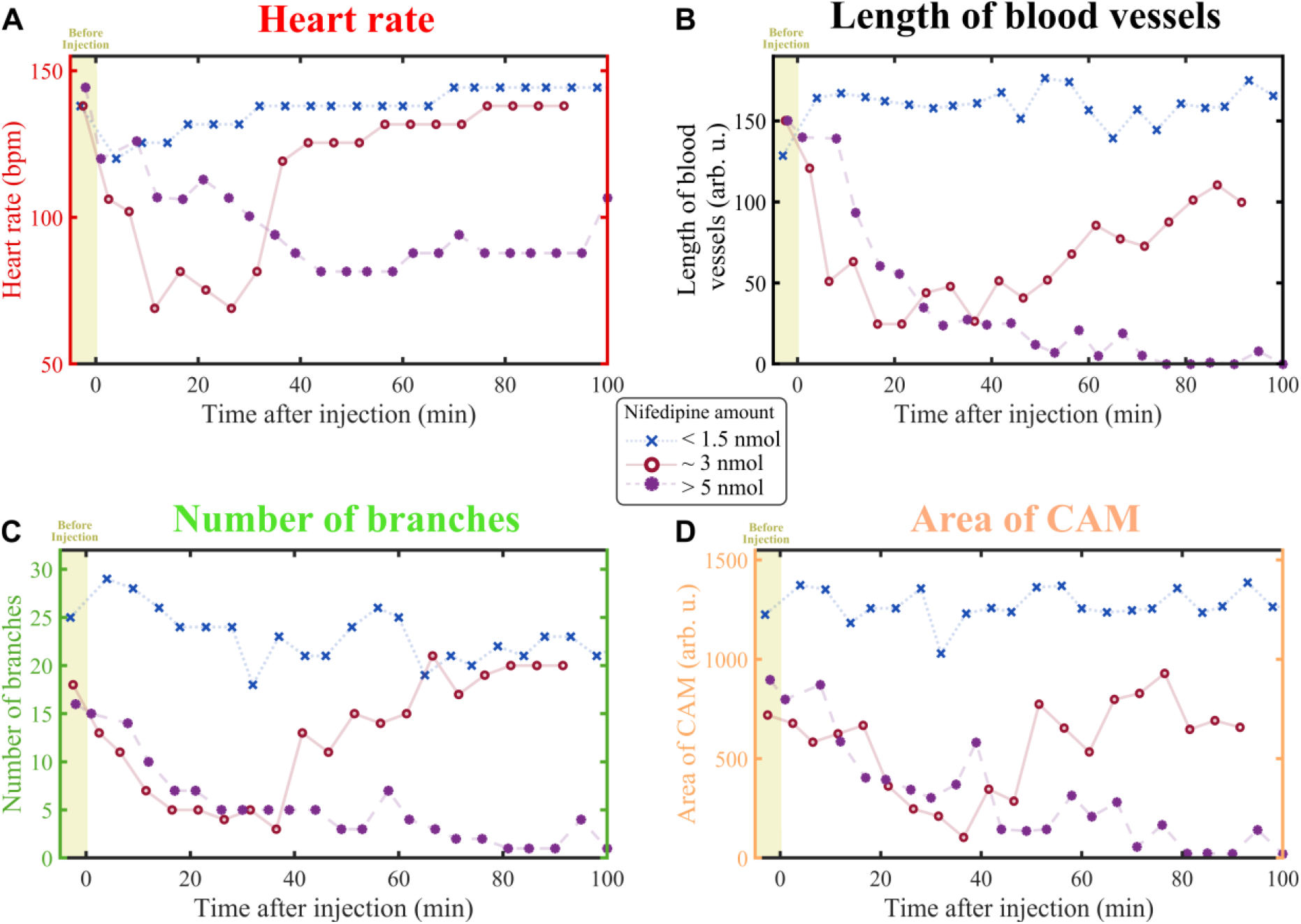
Impact of Nifedipine drug injection on the vascular structure of the chick embryo. The extracted features are: (A) Heart rate, (B) blood vessel length, (C) number of branches, and (D) CAM area at various times post-injection.

**Fig. 5.**
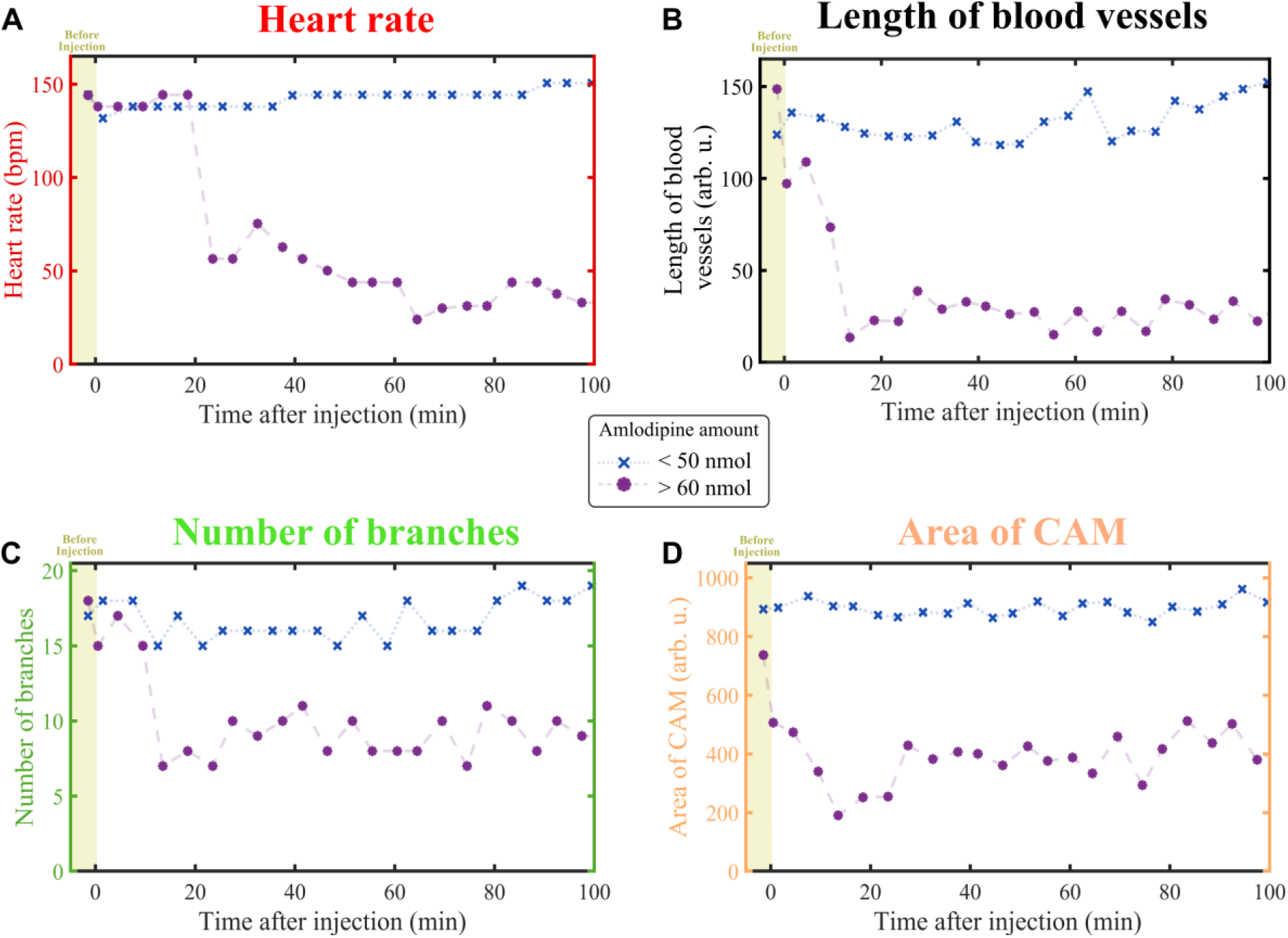
Impact of Amlodopine injection on the vascular structure of the chick embryo. The extracted features are: (A) Heart rate, (B) blood vessel length, (C) number of branches, and (D) CAM area at various times post-injection.

The chick embryo responded to the Nifedipine in a dose dependent manor (Fig.4 A-C). At the lowest dose tested (<1.5 nmol), apart from an initial drop during the drug injection in the bpm, the heart rate remained constant throughout the length of the experiment. The embryo developed normally when incubated for 7 days after the experiment. At low doses around 3 nmol, the heart rate dropped over a 40 minute period, after which it began increase reaching normal levels after an hour. At higher doses above 5 nmol, the heart rate of the embryo dropped rapidly reaching its nadir after 20 minutes. It did not begin to return normal levels even after 100 minutes after injection, indicating that the embryo is dying. At these higher doses the embryo died at the time of injection or its later development was heavily compromised. For each concentration, we repeated the tests on five different eggs, and similar behavior was observed.

### 3.4 Amlodipine injection into chicken eggs

The chick embryo also responded to Amlodipine in a dose dependent manor (Fig.5 A-C). Amlodipine was effective on the chick embryo at a dose that was ten times higher than that of Nifedipine. At the lowest dose tested (<50 nmol), apart from an initial drop during the drug injection in the bpm, the heart rate remained constant throughout the length of the experiement. The embryo also developed normally when incubated for 7 days after the experiment. At higher doses > 60 nmol, the heart rate of the embryo dropped rapidly reaching its nadir after 20 minutes. At these higher doses the embryo died at the time of injection or its later development was heavily compromised. For Amlodipine, we were unable to observe the intermediate case where an initial drop in heart rate and blood vessel circulation is followed by a revival. However, we do not rule out the possibility of such a scenario occurring with Amlodipine, but confirming this would require a more detailed study. For each concentration, we repeated the tests on five different eggs.

## 4. Discussion

We developed and applied an LSCI system for non-invasive, longitudinal tracking of chick embryo heart rate, blood flow, and blood vessel structure. The system is automated and does not alter the embryo’s natural development. We used this system to distinguish the effect between two closely related L-type Ca receptor antagonists Nifedipine and Amlodipine on chick embryos. Our findings indicate that Nifedipine is effective at reducing the heart rate and blood flow at concentration levels ten times lower than Amlodipine. Amlodipine was more stable and longer lasting. In control experiments, where only the solvent was injected, the heart rate and vascular structure remained consistent throughout the recording period. Although Nifedipine and Amlodipine belong to the same family of Ca+ channel antagonists, the LSCI system could distinguish between their effect onto the chick embryo.

Other imaging techniques, such as fluorescent microscopy or optical coherence tomography, have been effectively used to image live chick embryos and CAM vessels. However, these methods require a viewing window to be cut into the eggshell to reduce light scattering. In contrast, the LSCI system provides quantitative real-time imaging of the chick embryonic heart and extraembryonic blood vessels without breaking the shell. In our experiments, we used an infrared laser with an 852 nm wavelength and an illumination power of 220 mW. This laser power level did not compromise embryo development, as 7-day post-imaging embryos (at 12 days of age) displayed normal morphology after being imaged every 3 minutes over several hours and then incubated. Indeed, the eggshell acts as a protective layer by scattering and absorbing the light, ensuring that the amount of laser power reaching the embryo remains within a safe range.

All chick embryos imaged in this study were at stages HH16 to HH19. However, the range of imaging can be extended from stages HH11 to HH25, as our current LSCI system is capable of imaging embryos within this range. Imaging embryos beyond stage HH25 may not be feasible due to their larger body size, increased thickness, and body movements, which all degrade LSCI reconstruction. However, developmental stages before HH11 can be successfully imaged by adjusting the field-of-view, speckle size, and camera exposure time of the LSCI system, as long as the embryo exhibits blood circulation.

This new integrated LSCI system offers a rapid quantitative monitoring of the chick heart function and vascular structure. It can be used to test agents that affect the heart as well as the blood vessels. In the future it could be used for longitudinal studies of heart development to test factors such as retinoic acid (RA) which is crucial to cardiac development in all vertebrates (Perl and Waxman, 2020).

## Credit authorship contribution statement

**Carol Readhead:** Conceptualization, Formal analysis, Investigation, Methodology, Writing – original draft, Writing – review & editing,

**Simon Mahler**: Conceptualization, Formal analysis, Investigation, Methodology, Writing – original draft, Writing – review & editing,

**Zhenyu Dong:** Visualization, Methodology, Writing – original draft, Writing – review & editing,

**Yuki Sato:** Visualization and advice on injectio, Methodology, Writing – original draft, Writing – review & editing,

**Changhuei Yang:** Conceptualization, Superivision, Funding acquisition, Writing – original draft, Writing – review & editing,

**Marianne E. Bronner:** Conceptualization, Superivision, Funding acquisition, Writing – original draft, Writing– review & editing,

## Data availability

Data will be made available on request.

## Acknowledgements

The authors would like to thank Dr. Xi Chen for his help and advice on avian embryos.

